# Phosphorylation by CK2 Regulates MUS81/EME1 in Mitosis and After Replication Stress

**DOI:** 10.1101/278143

**Authors:** Anita Palma, Giusj Monia Pugliese, Ivana Murfuni, Veronica Marabitti, Eva Malacaria, Sara Rinalducci, Anna Minoprio, Massimo Sanchez, Filomena Mazzei, Lello Zolla, Annapaola Franchitto, Pietro Pichierri

## Abstract

The MUS81 complex is crucial for preserving genome stability through the resolution of branched DNA intermediates in mitosis. However, untimely activation of the MUS81 complex in S-phase is dangerous. Little is known about the regulation of the human MUS81 complex and how deregulated activation affects chromosome integrity. Here, we show that the CK2 kinase phosphorylates MUS81 at Serine 87 in late-G2/mitosis, and upon mild replication stress. Phosphorylated MUS81 interacts with SLX4, and this association promotes the function of the MUS81 complex. In line with a role in mitosis, phosphorylation at Serine 87 is suppressed in S-phase and is mainly detected in the MUS81 molecules associated with EME1. Loss of CK2-dependent MUS81 phosphorylation contributes modestly to chromosome integrity, however, expression of the phosphomimic form induces DSBs accumulation in S-phase, because of unscheduled targeting of HJ-like DNA intermediates, and generates a wide chromosome instability phenotype. Collectively, our findings describe a novel regulatory mechanism controlling the MUS81 complex function in human cells. Furthermore, they indicate that, genome stability depends mainly on the ability of cells to counteract targeting of branched intermediates by the MUS81/EME1 complex in S-phase, rather than on a correct MUS81 function in mitosis.

## INTRODUCTION

Resolution of branched DNA intermediates formed during recombination or upon replication stress is important for cell cycle progression and genome stability maintenance (1, 2). Consistently, cells have evolved multiple and redundant mechanisms to ensure processing of recombination/replication intermediates, minimising the risk of entering into mitosis and performing cell division with unreplicated or untangled chromosomes(3).

In eukaryotes, the Mus81/Mms4^EME1^ heterodimer, the Mus81 complex, is the major endonuclease involved in the resolution of recombination or replication DNA intermediates (4–6). The main physiological function of the Mus81 complex is performed during the G2/M phase of the cell cycle (1). However, recent evidence clearly show activation of the Mus81 complex also in S-phase under conditions of persisting replication stress (7–10). Interestingly, such “pathological” Mus81-dependent processing of replication intermediates would be essential for proliferation, but it is also involved in the generation of chromosome instability (7, 11–13). Hence, as unscheduled activation of endonucleolytic cleavage is detrimental for genome integrity, Mus81 complex activity needs to be tightly regulated. In yeast, regulation of the Mus81 complex is largely dependent on phosphorylation of the non-catalytic subunit Mms4 by the mitotic kinases Cdc28^PLK1^ and Cdc;(4, 14). Adding further complexity to the mechanism, activation of Mus81/Mms4 in mitosis has been recently shown to require Cdc7-Dbf4, another cell cycle-regulated kinase (15). In fission yeast, Mus81/Eme1 function is also positively controlled by Cdc2^CDK1^ and Chk1 in response to DNA damage while it is repressed by the checkpoint kinase Cds1^CHK2^, in S-phase (16, 17). Although evidence suggest that mitotic kinases may regulate the function of the MUS81 complex in human cells too, we do not have much mechanistic insights. The recent observation of the crucial role of CDK1-mediated phosphorylation of SLX4 in controlling MUS81 complex function does provide mechanistic clues, but also make difficult to distinguish between direct vs. indirect effects of CDK1 on MUS81-dependent resolution (18, 19). Moreover, although signs of phosphorylation-induced changes in the electrophoretic mobility of EME1 and EME2 are apparent (20), little if any information exist on phosphorylation of the MUS81 subunit and its possible functional relevance. However, phosphorylation of the invariant subunit of the two MUS81/EME complexes could be a more efficient way to regulate activity of the holoenzyme, as well as association with proteins that can influence its biological activity under normal or pathological conditions (1, 7, 20, 21). Hence, it is likely that the MUS81 undergoes cell-cycle-specific phosphorylation events that may contribute to tightly regulate its function together with the observed modification of the EME1/2 subunit.

In addition to PLK1 and CDK1, an attractive kinase for the regulation of MUS81 complex is the pleiotropic CK2 (22, 23). Indeed, CK2 is important to regulate mitotic progression, is activated by CDK1 and phosphorylates several repair/recombination enzymes, such as MDC1, MRE11 and RAD51 (22, 24–27).

Here, we found that CK2 phosphorylates the N-terminal region of MUS81 at Serine 87 (S87) in the early mitotic stage and after mild replication stress. Phosphorylation by CK2 stimulates MUS81/EME1 association with SLX4, and it is required for the biological function of the complex in mitosis. Introduction of a phosphomimetic mutation at S87 results in unscheduled function of MUS81 complex during DNA replication and accumulation of extensive genome instability already in untreated cells. Therefore, our results represent the first demonstration of regulatory phosphorylation of the MUS81 subunit of the MUS81 complex in human cells, which is specifically targeted at the MUS81/EME1 heterodimer and is crucial for its function during mitosis. As CK2 is upregulated in many tumours, deregulation of this mechanism may contribute to increase their genomic instability during cancer development.

## MATERIALS AND METHODS

### In vitro kinase assay

For in vitro kinase assays, 300ng or 2μg (MS/MS) of the indicated GST-fused MUS81 fragments were incubated with recombinant purified CK2 (NEB), kinase in the presence of ^32^ P-ATP, or ATP, and in kinase-specific reaction buffer prepared according to the manufacturers’ directions. After washing, GST-fragments were released and analysed as previously reported (28). Caseins (Sigma-Aldrich) were used as positive control in the CK2 kinase assay. For the full-length assay, 200ng of full-lenght MUS81 immunopurified from HEK293T cells (Origene) was incubated with 200ng of CK2 kinase. Phosphorylation was analysed by SDS-PAGE and WB.

### Cell culture, generation of cell lines and RNA interference

MRC5SV40 and HEK293T cell lines were maintained as described (28).

To obtain the MRC5shMUS81 cells or the shMUS81-HEK293T cells, a retroviral plasmid containing an MUS81-targeting shRNA sequence (Origene cod. TR303095, sequence #1) was nucleofected using the Neon system (Life technologies). Three days after nucleofection, cells were subjected to selection with 500ng/ml puromycin and resistant clones expanded, tested for MUS81 depletion and phenotype before further use. To complement MRC5 shMUS81 cells with the wild-type MUS81 or its phosphomutants, the wild type form of MUS81 ORF cloned into the pCMV-Tag2B plasmid was subjected to SDM (Quickchange II XL – Stratagene) to introduce the S87A or S87D mutations.

After the first round of mutagenesis, all MUS81 ORFs were made RNAi-resistant by SDM, sequence-verified and cloned into pEF1a-IRES-NEO vector by the Gibson Assembly protocol (NEB). Sequence-verified plasmids were then transfected into MRC5 shMUS81 cells by the Neon nucleofector (Life technologies) in order to obtain cell lines stably expressing MUS81 and its mutant forms. Cell colonies were selected by 1mg/ml G418 antibiotic (Santa Cruz Biotechnologies). RNA interference against MUS81 was performed as previously reported (11). In all experiments, cells were transfected using Lullaby (OZ Biosciences). RNA interference against EME2 was performed with siRNA-SMART pool, 10nmol (197342) from Dharmacon. The efficiency of protein depletion was monitored by western blotting 72h after transfection.

### Chemicals

Hydroxyurea (Sigma-Aldrich) was used at 2 mM, the DNA replication inhibitor aphidicolin (APH) (Sigma-Aldrich) was used at 0.2 μM (low dose) or 1.5 μM (intermediate dose). The CK2 inhibitor CX4925 (Selleck chemicals) was used at 25 μM, the CDK1 inhibitor (RO-3306, Sigma-Aldrich) at 9 μM and WEE1 inhibitor (MK-1775, Selleck chemicals) was used at 500 μM. Nocodazole (Sigma-Aldrich) was used at 0.5 μg/μl and Thymidine (Sigma-Aldrich) at 2 mM. 5-Bromo-2´-Deoxyuridine (BrdU, Sigma-Aldrich) was used at 30 μM.

### Neutral Comet assay

Neutral Comet assay was performed as previously described (28). A minimum of 200 cells was analyzed for each experimental point.

### Immunoprecipitation and Western blot analysis

Cell lysates and immunoprecipitation experiments were performed as previously described (Pichierri et al, 2012). FLAG-tagged proteins were immunoprecipitated with anti-FLAG M2 Agarose Beads (Sigma-Aldrich) while cMYC-tagged proteins were purified with Myc-TRAP MA magnetic agarose beads (Chromotek). Western blot was performed using standard methods. Blots were developed using Westernbright ECL (Advasta) according to the manufacturer’s instructions. Quantification was performed on scanned images of blots using Image Lab software.

### PLA (Proximity-Ligation Assay)

The *in-situ* proximity-ligation assay (PLA; mouse/rabbit red starter Duolink kit from Sigma-Aldrich) was used as indicated by the manufacturer. Images were acquired with Eclipse 80i Nikon Fluorescence Microscope, equipped with a VideoConfocal (ViCo) system. For each point, at least 500 nuclei were examined and foci were scored at 40×. Parallel samples incubated with only one primary antibody confirmed that the observed fluorescence was not attributable to artefacts. Only nuclei showing more than four bright foci were counted as positive.

### Antibodies

The primary antibodies used were: anti-MUS81 (1:1000; Santa Cruz Biotechnologies), anti-DDK (Flag Origene, WB 1:1000, IF 1:200), anti-cMYC (1:1000;Abcam),, anti-CK2 (1:1000; Cell signaling technologies), anti-RxxpS/T (1:1000; Cell signaling technologies), anti-SLX4 (WB 1:1000, IF 1:200, Novus biologicals), anti-pS10H3 (1:1000, Santa Cruz Biotechnologies), anti-H3(1:1000, Santa Cruz Biotechnologies), anti-EME1 (1:1000, Santa Cruz Biotechnologies), anti-EME2 (1:500, Invitrogen), anti-Cyclin A (WB: 1:1000, IF: 1:100, Santa Cruz Biotechnologies), anti 53BP1 (1:400, Millipore), anti-BrdU (1:50, Becton Dickinson), anti-pS139H2A.X (1:1000,Millipore), anti-ɣ-Tubulin (1:200, Sigma-Aldrich), anti-pS87MUS81 (WB 1:1000, IF 1:200, Abgent) and anti-Lamin B1 (1:10000; Abcam). HRP-conjugated matched secondary antibodies were from Jackson Immunoresearch and were used at 1:40000.

### Chromatin fractionation

Cells (4 × 10^6^cells/ml) were resuspended in buffer A (10 mM HEPES, [pH 7.9], 10 mM KCl, 1.5 mM MgCl2, 0.34 M sucrose, 10% glycerol, 1 mM DTT, 50 mM sodium fluoride, protease inhibitors [Roche]). Triton X-100 (0.1%) was added, and the cells were incubated for 5 min on ice. Nuclei were collected in pellet by low-speed centrifugation (4 min, 1,300 ×g, 4°C) and washed once in buffer A. Nuclei were then lysed in buffer B (3 mM EDTA, 0.2 mM EGTA, 1 mM DTT, protease inhibitors). Insoluble chromatin was collected by centrifugation (4 min, 1,700 × g, 4°C), washed once in buffer B + 50mM NaCl, and centrifuged again under the same conditions. The final chromatin pellet was resuspended in 2X Laemmli buffer and sonicated for 15 s in a Tekmar CV26 sonicator using a microtip at 25% amplitude.

### Immunofluorescence

Immunofluorescence microscopy was performed on cells grown on cover-slips as described previously (28). Nocodazole-treated cells were blocked and fix with PTEMF buffer (29). After blocking, coverslips were incubated for 1hat RT with the indicated antibodies. For detection of anti-BrdU, after permeabilization with 0,4%Triton-X 100/PBS, cells were denatured in HCl 2,5N for 45’ at RT. Alexa Fluor^®^ 488 conjugated-goat anti mouse and Alexa Fluor^®^ 594 conjugated-goat anti-rabbit secondary antibodies (Life Technologies) were used at 1:200. Nuclei were stained with 4’,6-diamidino-2-phenylindole (DAPI 1:4000, Serva). Coverslips were observed at 40× objective with the Eclipse 80i Nikon Fluorescence Microscope, equipped with a VideoConfocal (ViCo) system. Images were processed by using Photoshop (Adobe) program to adjust contrast and brightness. For each time point at least 200 nuclei were examined. Parallel samples incubated with either the appropriate normal serum or only with the secondary antibody confirmed that the observed fluorescence pattern was not attributable to artefacts.

Experiments for labeling cellular DNA with EdU (5-ethynyl-2’-deoxyuridine). EdU was added to the culture media (10μM), for 30 min. Detection of EdU was performed used Click-iT EdU imaging Kits (Invitrogen).

### Cell cycle analysis by flow cytometry

Cells were processed for flow cytometry as follows: for each point, 10^6^ cells were collected, and after two washes in PBS, fixed in 70% cold ethanol. Then, cells were washed in PBS/BSA 1% and then resuspended in 0.5μg/ml propidium iodide and 0.1mg/ml RNase before analysis. Data were analysed with CellQuest and ModFit LT 4.1. software.

Bivariate flow cytometry was performed for anti-BrdU and anti-γ-H2AX staining as indicated in the Anti-BrdU data-sheet (Becton Dickinson).

### Growth Curve

The cells were seeded at 1.8 × 10^4^ cells per plate. After trypsinization, cells were counted through electronic counting cells (BioRad) for the following 6 days. The growth curve of the cell cultures was expressed as number of cells as a function of time.

### Chromosomal aberrations

Cells for metaphase preparations were collected according to standard procedure and as previously reported (30). Cell suspension was dropped onto cold, wet slides to make chromosome preparations. The slides were air dried overnight, then for each condition of treatment, the number of breaks and gaps was observed on Giemsa-stained metaphases. For each time point, at least 50 chromosomes were examined by two independent investigators and chromosomal damage was scored at 100x; magnification with an Olympus fluorescence microscope.

### Statistical analysis

All the data are presented as means of at least three independent experiments. Statistical comparisons were made by Student’s *t* test or by Anova, as indicated. P < 0.5 was considered significant.

## RESULTS

### MUS81 is phosphorylated at Serine 87 by the protein kinase CK2 both *in vitro* and *in vivo*

The human MUS81 contains an N-terminal unstructured region and a HhH domain that are essential to associate with SLX4, a crucial step for the biological function of the complex in mitosis (31). Hence, seeking for regulatory events modulating the MUS81 complex, we scanned the N-terminal sequence of MUS81 comprising amino acids 1-200, which includes the SLX4 binding region, for the presence of putative phosphorylation sites of mitotic kinases. As shown in Table 1, bioinformatics analysis retrieved several CDKs, PLK1 and CK2 putative phosphorylation sites that score over the specificity threshold of the software. As CK2 was not previously associated to MUS81 regulation and its pharmacological inhibition interfered with the formation of MUS81-dependent DSBs in checkpoint-deficient cells (7, 32) (Fig. S1), we decided to focus on this kinase. Hence, to test whether CK2 could phosphorylate MUS81 *in vitro,* we performed a radioactive kinase assay. As substrates, we used two different fragments comprising residues 1-206 or 76-206 of MUS81, fused to GST and purified from bacteria (Fig. 1 A). Our kinase assays showed that CK2 efficiently phosphorylates the N-terminal MUS81 fragments (Figs. 1 B).

**Table 1.**
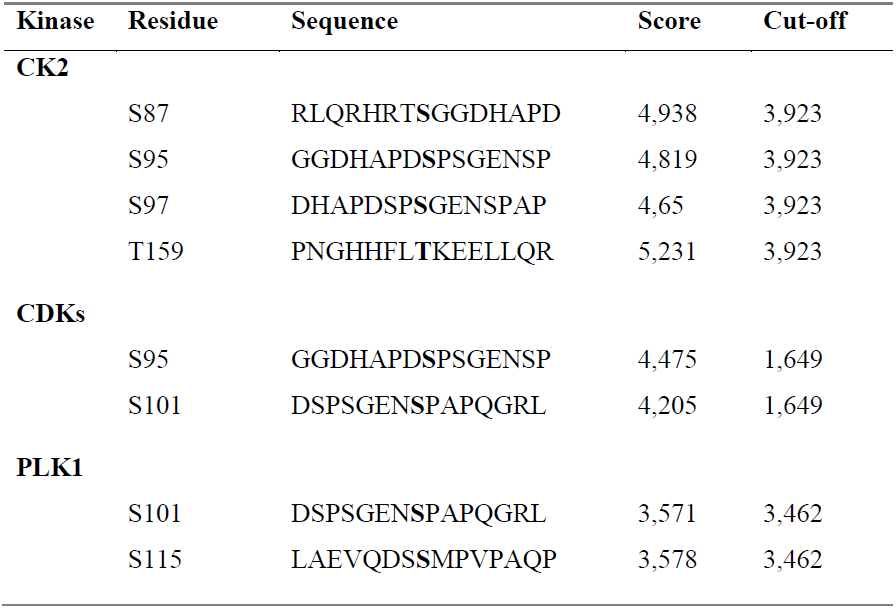
Putative phosphorylation sites identified in the N-terminal MUS81 region

| **Kinase** | **Residue** | **Sequence** | **Score** | **Cut-off** |
| --- | --- | --- | --- | --- |
| **CK2** |  |  |  |  |
|  | S87 | RLQRHRTSGGDHAPD | 4,938 | 3,923 |
|  | S95 | GGDHAPDSPSGENSP | 4,819 | 3,923 |
|  | S97 | DHAPDSPSGENSPAP | 4,65 | 3,923 |
|  | T159 | PNGHHFLTKEELLQR | 5,231 | 3,923 |
| **CDKs** |  |  |  |  |
|  | S95 | GGDHAPDSPSGENSP | 4,475 | 1,649 |
|  | S101 | DSPSGENSPAPQGRL | 4,205 | 1,649 |
| **PLK1** |  |  |  |  |
|  | S101 | DSPSGENSPAPQGRL | 3,571 | 3,462 |
|  | S115 | LAEVQDSSMPVPAQP | 3,578 | 3,462 |

**Figure 1.**
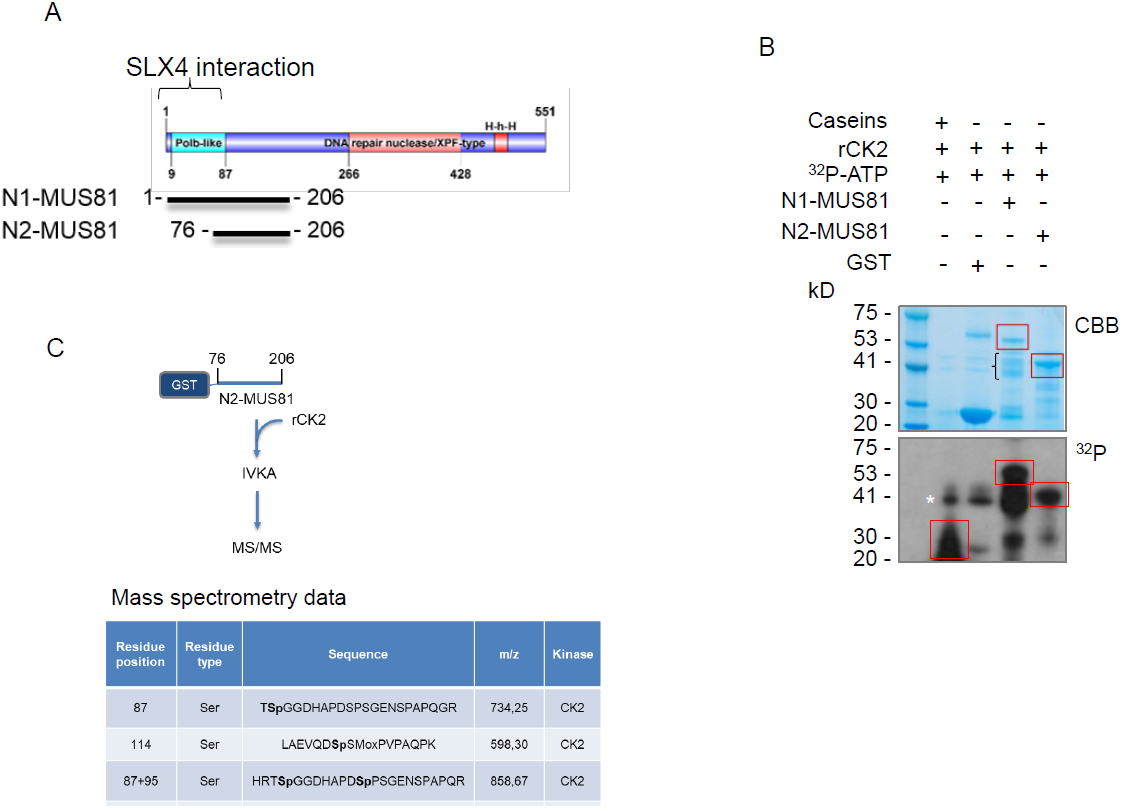
CK2 phosphorylates MUS81 *in vitro.* (A) Schematic representation of the N-terminal MUS81 fragments. (B) GST-fused N-terminal MUS81 fragments purified from bacteria were incubated with recombinant CK2 and subjected to in vitro radioactive kinase assay. Caseins were used as positive control. Red boxes indicate position of the fragments. Parenthesis indicates N1-MUS81 degradation products. Asterisk denotes residual co-purifying autophosphorylated CK2. (C) MS/MS analyses of in vitro phosphorylated N2-MUS81. Inset summarizes the identified phosphopeptides and residues.

To confirm phosphorylation and identify phosphoresidues, we incubated the fragment 76-206 of MUS81 with recombinant CK2 and analysed the product of the reaction by MS/MS after affinity-purification of the phosphorylated peptides. From the CK2-modified fragment, we identified three different peptides containing phosphorylated residues (Fig. 1 C). Among the three identified residues, S87 was the most promising because it is very close to the SLX4-interacting region of MUS81 (33). Hence, to functionally characterize this phosphorylation event, we generated a phosphospecific antibody that recognizes the MUS81 protein modified at S87. Dot blot assay confirmed that the anti-pS87MUS81 antibody efficiently recognizes the modified peptide or the peptide incubated with recombinant CK2, while it showed no antibody reaction with the peptide incubated with recombinant PLK1 (Fig. S2 A). Phosphorylation of MUS81 was also confirmed using a commercial phosphomotif antibody that recognizes a sequence (**R**XX**pS/T**) very similar to that surrounding S87 (**R**HRT**pS**) (Fig. S2 B). Finally, co-immunoprecipitation experiments revealed interaction between the catalytic subunit of the CK2 holoenzyme, CK2α, and MUS81, supporting the possible physiological relevance of S87 modification (Fig. S2 C).

Next, we performed *in vivo* experiments to test the ability of the anti-pS87MUS81 antibody to detect MUS81 phosphorylation. To this aim, HEK293T cells stably expressing a shRNA sequence against MUS81 (HEK293TshMUS81) were transiently transfected with empty vector or with plasmids expressing the wild-type, the unphosphorylable (S87A) or the phosphomimetic (S87D) FLAG-tagged RNAi-resistant form of MUS81 protein. After anti-FLAG immunoprecipitation, the presence of MUS81 phosphorylation was analysed by Western blotting. As shown in Fig. S3 A, S87 phosphorylation was detected only in the wild-type protein, confirming S87 modification *in vivo* and the specificity of the antibody. This result was further corroborated by immunofluorescence in MRC5SV40 cells stably expressing the shMUS81 construct (shMUS81), and shMUS81 cells complemented with the FLAG-tagged-RNAi-resistant form of wild-type MUS81 or each of the two phosphorylation mutants and enriched in mitosis using nocodazole (Fig. S3 B). Our analysis revealed the presence of nuclear staining in MUS81^WT^ cells, which was not detectable in shMUS81 cells or in cells expressing the MUS81 mutant forms (MUS81^S87A^ and MUS81^S87D^ Fig. S3 B). Similarly, evaluation of MUS81 S87 phosphorylation by Western blotting in transiently expressing cells enriched in M-phase by nocodazole treatment confirmed that the anti-pS87 antibody efficiently recognised the MUS81 wild-type but not the phosphorylation mutant forms (Fig. S3 C). Interestingly, pharmacological inhibition of CK2 substantially reduced S87 phosphorylation as evaluated by IP/WB or immunofluorescence analysis (Figs. S4 A and B), proving that S87 residue of MUS81 is an *in vivo* substrate of the protein kinase CK2.

Altogether, our findings show that MUS81 is phosphorylated by CK2 on S87 both *in vitro* and *in vivo*, and that S87 phosphorylation is already detectable during unperturbed cell growth.

### CK2-mediated phosphorylation of MUS81 at Serine 87 is an early mitotic event stimulated by mild replication stress and restrained in the MUS81/EME2 complex

The MUS81/EME1 complex functions primarily in G2/M, while the MUS81/EME2 in S-phase (4, 34). Hence, we analysed if modification by CK2 was cell cycle-dependent. To this aim, HEK293TshMUS81 cells, transiently expressing the wild-type form of MUS81, were synchronized and S87 phosphorylation was determined. Phosphorylation was evaluated by IP/WB using the anti-pS87MUS81 antibody from cells enriched in S-phase after release from a double-thymidine block or in mitosis using Nocodazole (Noc; Fig. 2 A). Although MUS81 was found phosphorylated at S87 already in asynchronous cells, the level of phosphorylation was substantially reduced in S-phase enriched cells, but it was increased in mitotic cells (Fig. 2 B). To evaluate phosphorylation in a more physiological context, we performed anti-pS87MUS81 immunofluorescence in asynchronous cultures exposed to a short EdU pulse to label S-phase cells or, as a control, in Noc-arrested cultures. Dual EdU/pS87MUS81 immunostaining revealed that CK2-dependent phosphorylation is absent in S-phase cells, while it is easily detected after Noc treatment (Fig.2C).

**Figure 2.**
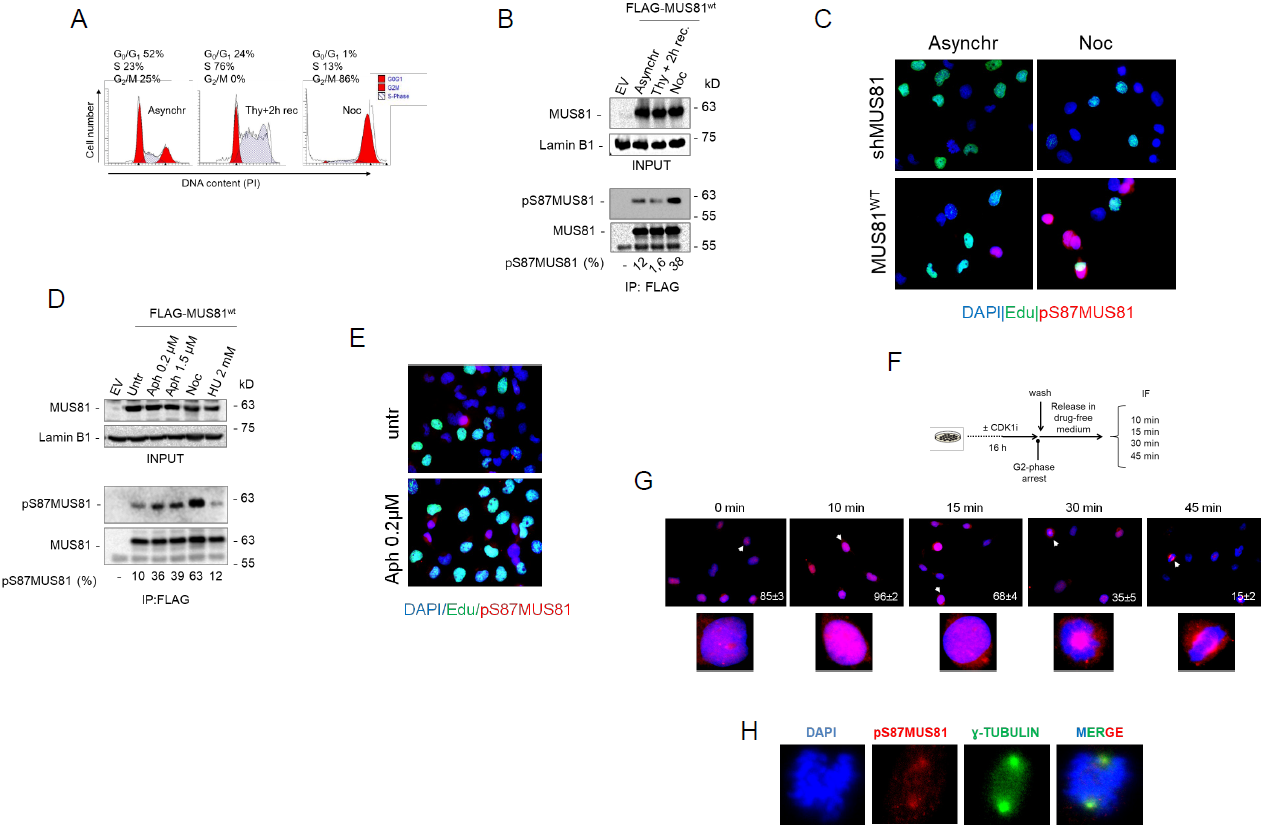
Phosphorylation of MUS81 on S87 is cell cycle-dependent and confined in early mitosis. (A) Flow cytometry analysis of HEK293T shMUS81 cells transiently expressing the wild-type form of FLAG-MUS81 after synchronization in S-phase (Thy+2h rec) or in mitosis (Noc). (B) After transient transfection, FLAG-MUS81^wt^ was immunoprecipitated from asynchronous (asynchr), S-phase or M-phase synchronized HEK293T shMUS81. Phosphorylation was analyzed by WB by using the anti-pS87MUS81 antibody, and the anti-pS87MUS81 signal was expressed as normalized percentage of the total immunoprecipitated MUS81. EV=Empty Vector (C) Anti-pS87MUS81 immunofluorescence staining (red) was performed in MRC5SV40 shMUS81 and FLAG-MUS81^wt^ stably complemented cells previously exposed to short EdU pulse to mark S-phase cells (green). Nuclei were depicted by DAPI staining (blue). (D) After transient transfection, FLAG-MUS81^wt^ was immunoprecipitated from asynchronous HEK293T shMUS81 cells treated as indicated. anti-pS87MUS81 signal was expressed as normalized percentage of the total immunoprecipitated MUS81. EV=Empty Vector. (E) Immunofluorescence experiments were performed to detect S87MUS81-phosphorylation upon mild replication stress induced by low dose of Aphidicolin. MRC5 shMUS81 cells complemented with MUS81wt were exposed to short EdU pulse and then stained with anti-pS87 antibody (red) and DAPI (blue). Representative images were shown. (F) Experimental scheme used to study S87 MUS81 phosphorylation over time, from G2-phase arrested cells to late mitosis. MRC5 shMUS81 cells complemented with MUS81^wt^ were arrested in G2-phase by treatment with CDKi (RO-3306). Cells were immunostained for anti-pS87MUS81 at indicated post-release time points. (G) Representative images of anti-pS87MUS81 immunostaining (red). The mean percentage of pS87MUS81 positive cells is indicated in the images ± SE. At least 200 anaphase, 100 metaphase and 50 prophase cells were analysed in three replicates. Arrowheads indicate the positive cells enlarged in the relative insets. (H) Representative images of anti-pS87 MUS81 antibody (red) co-localization with γ-Tubulin antibody (green) in metaphase cells. Nuclei were depicted with DAPI.

Although the most relevant function of the MUS81 complex is the resolution of recombination intermediates in late G2/M, it may also process perturbed or collapsed replication forks (1). Thus, we analysed if phosphorylation of S87 might be a common readout of the MUS81 complex activation. To this end, we treated HEK293TshMUS81 cells expressing the wild-type FLAG-MUS81 protein with two doses of aphidicolin (Aph), which partially arrest replication or perturb common fragile sites (CFS), or with hydroxyurea (HU). Phosphorylation was then assessed in anti-FLAG IP by WB using the anti-pS87MUS81 antibody. All these treatments have been reported to stimulate the function of both the MUS81 complexes, however, prolonged treatment with HU leads to a complete arrest of S-phase progression and formation of MUS81-dependent DSBs (8, 12, 13, 35). Consistently, Aph treatment accumulated cells in S-phase but did not completely arrest cell cycle progression, while 24 h of HU blocked cells in G1/S phase (Fig. S5). As expected, phosphorylation of S87 was increased in Noc-treated cells, but it was also stimulated by Aph treatments (Fig. 2 D). In contrast, and despite the reported formation of DSBs by MUS81, phosphorylation of S87 was barely detectable in cells treated with HU (Fig. 2 D). To confirm that treatment with Aph stimulated phosphorylation of MUS81 at S87, we performed anti-pS87MUS81/EdU immunofluorescence in shMUS81 cells complemented with the wild-type form of MUS81 (Fig. 2 E). Treatment with a low-dose Aph increased the number of nuclei positive to anti-pS87MUS81 immunostaining, although the large majority of cells staining positive for pS87 were EdU-negative. This confirms that a mild replication stress associated with activation of the MUS81 complex in late G2/M phase induced a CK2-dependent phosphorylation of MUS81.

Although it is widely accepted that the MUS81 complex carries out its primary function in late G2 and mitosis, it is still unclear if it is active throughout all this period. Our data suggest that phosphorylation at S87 can be used as diagnostic sign of MUS81 complex function. Hence, we performed immunofluorescence to follow MUS81 modification over time after release from a G2-arrest induced by CDK1 inhibition (29), as outlined in the experimental scheme (Fig. 2 E). Cells released in late G2/M were subjected to anti-pS87MUS81 immunofluorescence at different time-points. Cells blocked in late-G2 showed high levels of pS87MUS81 immunostaining, which increased during the early time-point after release in mitosis (Fig. 3 B). Anti-pS87MUS81 nuclear staining declined thereafter in concomitance with appearance of metaphase cells, as evaluated by DAPI staining (Fig. 2 F). Interestingly, pS87MUS81 nuclear immunostaining was always confined to cells with morphological features of late G2/prophase, even at later post-release time-points, when population was enriched of metaphase and anaphase cells (30 and 45min; Fig. 2 F). Of note, in several late mitotic cells, pS87MUS81 immunostaining was apparently accumulated at centrosomal regions (see 45min) as indicated by anti-γ-tubulin co-staining (Fig.2G).

**Figure 3.**
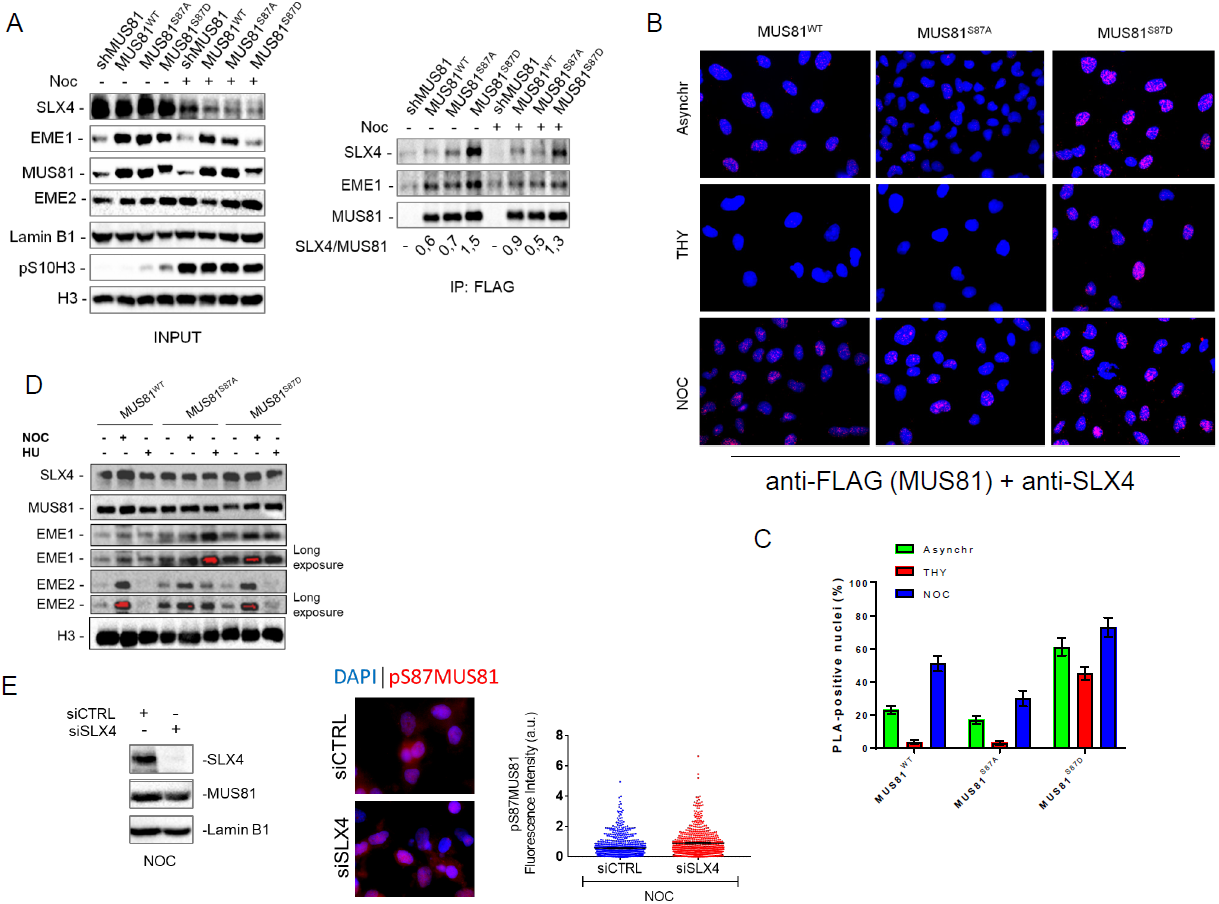
S87-MUS81 phosphorylation regulates SLX4 binding. (A) MUS81 was immunoprecipitated from asynchronous or M-phase synchronized cells expressing MUS81^wt^ or its phosphorylation mutants. The fraction of SLX4 associated to MUS81 was normalized on the total immunoprecipitated MUS81. (B) Interaction of MUS81 and SLX4 was analysed by PLA in asynchronous, THY and NOC-synchronized cells. Representative images of PLA fields are shown. MUS81-SLX4 interaction resulted in red nuclear dots. The mean percentage of PLA-positive cells is represented in the graph in (C). (D) Chromatin fraction was prepared from cells expressing MUS81^wt^ or its phosphorylation mutants, and treated as indicated. The presence of MUS81 and the indicated proteins in chromatin was assessed by WB. Histone H3 was used as loading control protein. Blots are representative of two different biological replicates, (E) MUS81 phosphorylation was evaluated by anti-pS87MUS81 immunofluorescence on cells transfected siCTRL or siSLX4 and accumulated in M-phase with Noc. The WB shows actual depletion levels, and images are representative of IF. The dispersion graph shows quantification of pS87MUS81 antibody signal intensities.

In human cells, MUS81 exists as an heteroduplex in association with EME1 or EME2 (36). Although there are conflicting results about the cell cycle-dependent association with EME2 (18, 20), EME1 is found throughout the cell cycle, even if its function predominates in G2/M (20). Since phosphorylation of S87 is stimulated in early mitosis and is absent in S-phase synchronized cells, we investigated if it was confined to the MUS81/EME1 complex. To this end, we transiently expressed FLAG-MUS81 and Myc-EME2 or Myc-EME1 in HEK293T cells, and immunopurified the fraction of MUS81 associated with EME2 or EME1 by anti-Myc immunoprecipitation. We analysed S87 MUS81 phosphorylation in asynchronous cells or in cells accumulated in M-phase with Noc. As shown in Fig. S6, S87 MUS81 phosphorylation was detectable in both the MUS81 complexes but with different levels. In the MUS81/EME1 complex, S87 MUS81 phosphorylation was enhanced by nocodazole treatment by about 3-fold over the basal level while, in the MUS81/EME2 complex phosphorylation was similar in asynchronous cells but did not change after nocodazole treatment.

Collectively, these results demonstrate that CK2 phosphorylates MUS81 at S87 in early mitosis, and that this phosphorylation is stimulated in the presence of mild replication stress. Furthermore, they suggest that phosphorylation of S87 is prevented in S-phase and mainly concerns the MUS81/EME1 complex.

### Phosphorylation of MUS81 at Serine 87 by CK2 regulates binding to SLX4

Association of the MUS81 complex with SLX4 is essential for its function (31), and given that the SLX4-binding region is close to S87, we asked whether phosphorylation could influence this interaction. To this aim, we performed co-immunoprecipitation experiments in shMUS81 cells complemented with the wild-type form of wild-type MUS81 or each of the two phosphorylation mutants. In asynchronous wild-type or MUS81^S87A^ cells, little SLX4 was found in a complex with MUS81 (Fig. 3 A). However, a 2-fold higher amount of SLX4 co-immunoprecipitated with MUS81 in MUS81^S87D^ cells (Fig. 3 A). In Noc-treated cells, association of MUS81 with SLX4 was more affected by loss of S87 phosphorylation. Indeed, the unphosphorylable MUS81 mutant (MUS81^S87A^) co-immunoprecipitated less SLX4 than the wild-type or the phosphomimic form (MUS81^S87D^; Fig. 3 A). Interestingly, while the amount of SLX4 associated with MUS81 was increased by Noc treatment in wild-type cells, any modulation was detected in the MUS81^S87A^ or in the MUS81^S87D^ mutant (Fig. 3 A). In contrast, MUS81 immunoprecipitated similar amounts of EME1 independently on the phosphorylation status of S87 (Fig. 3 A). To further confirm that phosphorylation of S87 affected the interaction of MUS81 with SLX4, we analysed protein-protein interaction at the single cell level by the Proximity-Ligation Assay (PLA). Interestingly, and consistently with biochemical assays, association of MUS81 with SLX4 was enhanced in MUS81^S87D^ cells, while it was only barely detectable in wild-type cells or in cells expressing the unphosphorylable protein, as evaluated by the number of PLA-positive cells (Fig. 3 B and C). Of note, association of MUS81 with SLX4 is strongly enhanced in the presence of the phosphomimic mutant (MUS81^S87D^) also in cells enriched in S-phase by a thymidine block (Fig. 3 B and C), even if their association should be actively prevented at this stage to avoid targeting of replication intermediates (18). To test if enhanced association between MUS81 and SLX4 might correlate with an increased association with chromatin, we performed cellular fractionation experiments in cells treated with Noc or exposed to HU for 24 h, a condition in which we observed little if any S87 phosphorylation (Fig. 2 D). Western blotting analysis of the chromatin fraction with an anti-MUS81 antibody showed no substantial difference in the two phosphorylation mutants, as compared to the wild-type (Fig. 3 D). In a similar way, the fraction of chromatin-associated SLX4 did not vary among the different MUS81 forms, however, the amount of EME1 in chromatin was substantially elevated in asynchronous or Noc-arrested cells expressing the phosphomimetic mutant (Fig. 3 D). In contrast, expression of the unphosphorylable MUS81 mutant enhanced the level of chromatin-associated EME1 and EME2 in cells treated with 24h HU, and also increased the amount of EME2 in asynchronous, untreated, cells (Fig. 3 D).

As our results indicate that a mutation mimicking constitutive S87 MUS81 phosphorylation stimulates association with SLX4 but not chromatin recruitment, we decided to analyse if this interaction might be required for subsequent phosphorylation of MUS81 by CK2. Hence, we analysed S87 phosphorylation by anti-pS87MUS81 IF in cells transfected or not with siRNAs against SLX4 and accumulated in mitosis with Noc. Depletion of SLX4 did not prevent S87 MUS81 phosphorylation, which is indistinguishable from wild-type cells (Fig. 3 E).

Therefore, our data indicate that phosphorylation of S87 by CK2 is important to stabilize or stimulate the MUS81-SLX4 interaction. Moreover, they suggest that phosphorylation takes place before formation of the MUS81/EME1/SLX4 complex.

### Phosphorylation status of MUS81 at S87 controls unscheduled targeting of HJ-like intermediates in S-phase

By regulating interaction with SLX4, phosphorylation of MUS81 S87 by CK2 may have functional implications. Hence, we used cells expressing the S87 MUS81 phosphomutants as a very specific tool to analyse the phenotypic consequences of a deregulated MUS81-EME1 function, bypassing the need to interfere with cell-cycle kinases. Our analysis of cell growth evidenced a substantial delay in the proliferation of MUS81^S87D^ cells, while cells expressing the related unphosphorylable MUS81 mutant did not show any apparent defect as compared to the wild-type (Fig. 4 A). Delayed proliferation rate of MUS81^S87D^ cells did not correlate with substantial cell cycle defects (Fig. S7 A). Thus, we analysed if it could derive from accumulation of spontaneous DNA damage by doing immunofluorescence against γ-H2AX or 53BP1. MUS81 deficiency or expression of wild-type MUS81 resulted in low level of γ-H2AX-positive cells (< 2%) (Fig. 4 B). Very few nuclei were positive for γ-H2AX also in MUS81^S87A^ cells, however, their number was increased of about 10-fold in the phosphomimic S87D mutant (Fig. 4 B). Interestingly, in the S87D MUS81 mutant, almost the totality of the γ-H2AX-positive cells were also Cyclin A-positive (i.e. in S or G2 phase) and the large part (60%) were in S-phase (EdU-positive) (Figs. 4 C and D). Similarly, the portion of 53BP1-foci/Cyclin A double-positive cells in the population was increased by expression of the phosphomimic MUS81 mutant as compared to MUS81^S87A^ or wild-type cells (Fig. 4 E). Flow cytometry analysis of the γ-H2AX-positive population confirmed that DNA damage arises mostly in S-phase or in G2/M cells (Fig. S7 B). The higher load of DNA damage in cells expressing the phosphomimic S87D MUS81 protein prompted us to analyse if this could derive from unscheduled targeting of intermediates during DNA replication. We recently demonstrated that ectopic expression of a GFP-RuvA fusion protein is sufficient to interfere with formation of DSBs by structure-specific endonucleases in human cells during S-phase (37). Hence, we ectopically expressed GFP-RuvA in wild-type or in MUS81^S87D^ cells and analysed the presence of DNA damage by neutral Comet assay. As shown in Fig. 5 A, the amount of spontaneous DSBs detected in wild-type cells did not decrease upon expression of RuvA. However, expression of RuvA significantly decreased the accumulation of DSBs in MUS81^S87D^ cells. Similarly, when we analysed the formation of 53BP1 foci after ectopic RuvA expression, we observed a substantial reduction of the 53BP1 focus-forming activity especially in cells expressing the S87D MUS81 mutant (Fig. 5 B and C). To determine if the incidental cleavage of HJ-like intermediates triggered by deregulated phosphorylation of MUS81 at S87 might correlate also with increased enzymatic activity, we immunopurified FLAG-MUS81 complexes from cells transiently over-expressing the wild-type form of MUS81 or its phosphorylation mutants and assessed the associated endonuclease activity after incubation of anti-FLAG immunoprecipitates with a model nicked HJ, one of the preferred *in vitro* MUS81 substrates (Fig. S8 A). As shown in Fig. S8 B, wild type MUS81 cleaved the substrate with comparable efficiency between asynchronous and M-phase enriched cells, while the MUS81-S87D mutant increased its activity in nocodazole-treated cells. In contrast, the unphosphorylable MUS81 mutant showed always very little activity. Of note, and in agreement with our PLA and the CoIP data (see Fig. 3), less SLX4 was found associated with the unphosphorylable MUS81 mutant (Fig. S8C). Surprisingly, loss of S87 phosphorylation MUS81 increased the ability of the protein to immunoprecipitate EME2 although the amount of EME1 was unchanged.

**Figure 4.**
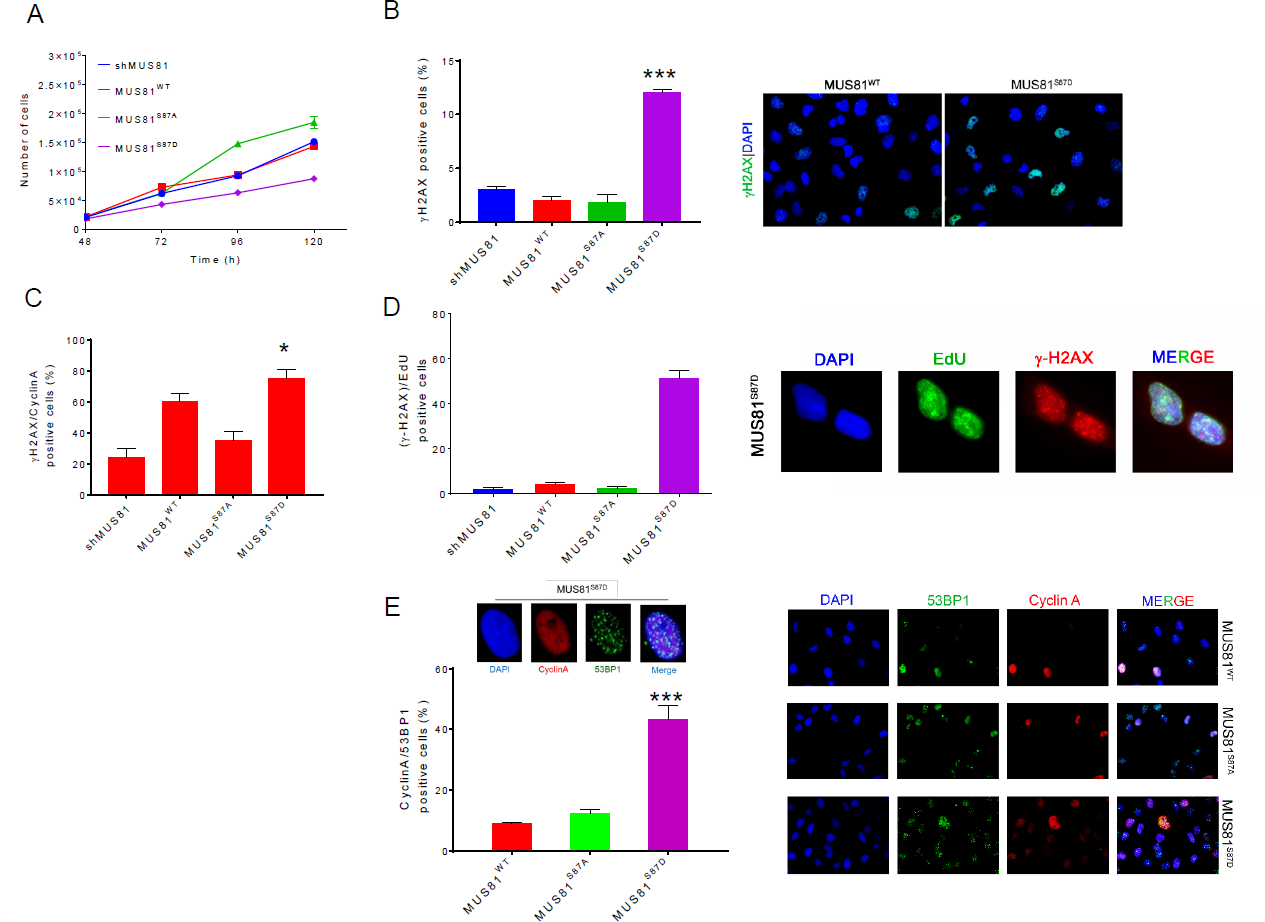
Constitutive phosphorylation of MUS81 at S87 affects cell growth and induces DNA damage. (A) Growth curve in shMUS81 cells stably expressing the wild-type form of MUS81 and it S87 phosphorylation mutants. Each point represents the average number of counts from two independent experiments. (B) Accumulation of spontaneous damage in MUS81 phosphorylation mutants. Untreated cells were immunostained with γ-H2AX antibody, the graph represents the analysis of γ-H2AX positive cells. Representative images of fluorescence fields are shown (γ-H2AX antibody: green; nuclear DNA: blue). (C) DNA damage, in S/G2-phase cells, was detected by double IF using anti-Cyclin A and γ-H2AX antibodies. (D) Analysis of the level of DNA damage during replication. Cells in S-phase were labelled by an EdU pulse and the graph shows the percentage of EdU (green) and anti-γ-H2AX (red) positive cells. Nuclear DNA was counterstained by DAPI (blue). (E) Analysis of 53BP1 foci formation in Cyclin A positive cells. Representative image of fluorescence cells stained with anti-53BP1 (green) and anti-Cyclin A (red) antibodies. Nuclear DNA was counterstained by DAPI. The graph shows quantification of the 53BP1-positive Cyclin A-cells. Data are mean values from three independent experiments. Statistical analysis was performed by Student’s t-test. Significance is reported compared to the wild-type: *** = p < 0.01; *= p< 0.05.

**Figure 5.**
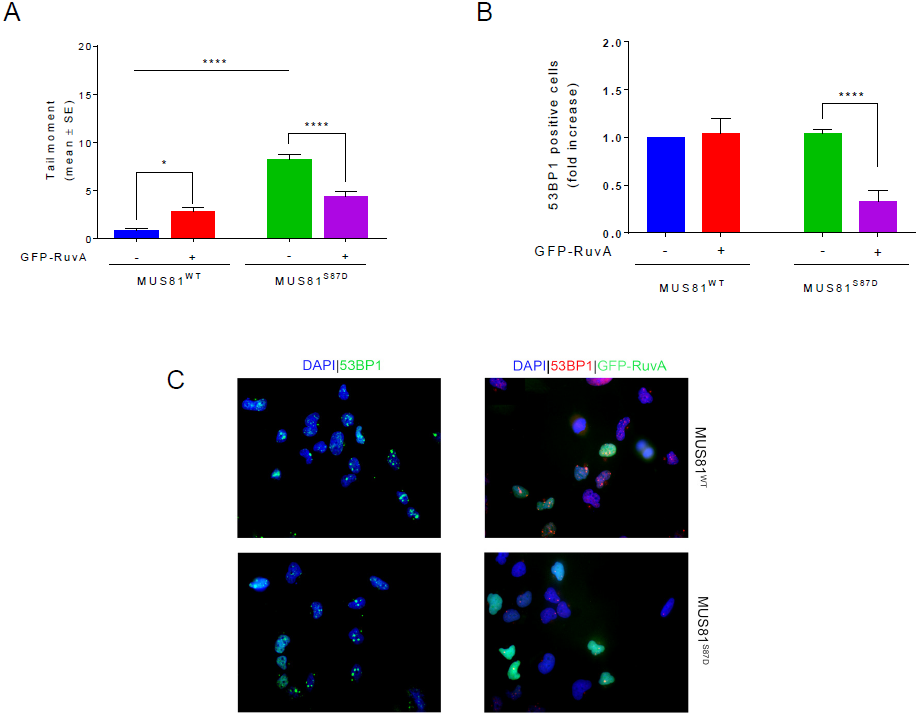
Ectopic GFP-RuvA expression reduces formation of DSBs and 53BP1 foci in constitutive-active MUS81 mutant. (A) MRC5 shMUS81 cells, stably expressing WT or S87D phosphomimetic MUS81 were transfected or not with GFP-RuvA. DSBs were evaluated 48h after transfection by neutral Comet assay. (B) The presence of 53BP1 nuclear foci was evaluated by IF 48h after transfection. The graph shows the fold increase of 53BP1 foci-positive cells over the untransfected cells. <u>Data are presented as mean ± standard error (SE) from three independent</u> <u>experiments. * = p < 0.5; **** = p< 0.001, ANOVA test.</u> (C)Representative microscopy fields are presented: 53BP1 antibody (green or red), GFP-RuvA and nuclear DNA was counterstained by DAPI (blue).

Collectively our findings indicate that deregulated MUS81 phosphorylation at S87 and subsequent MUS81-EME1-SLX4 association is sufficient to induce DNA damage in S-phase because of incidental cleavage of HJ-like intermediates, which is not correlated with enhanced enzymatic activity.

### Phosphorylation at Serine 87 induces premature mitotic entry through unscheduled MUS81 function in S-phase

Recent data evidenced that inhibition of WEE1 leads to premature activation of the MUS81 complex, because it releases CDK1 inhibition and stimulates phosphorylation of SLX4, licensing the formation of the MUS81/SLX4 complex in S-phase, and inducing uncontrolled progression in mitosis of unreplicated cells (18). We show that phosphorylation of MUS81 at S87 is crucial to MUS81-EME1 activation. Hence, we analysed if abrogation of S87 phosphorylation could revert the effect of WEE1 inhibition. To analyse premature mitotic entry from S-phase, we pulse-labelled a subset of replicating cells with BrdU, chased them in BrdU-free medium in the presence or absence of WEE1 inhibitor, and analysed progression of the BrdU-positive population through the cell cycle by bivariate flow cytometry (Fig. 6 A). Under unchallenged conditions, no substantial difference in the progression of the labelled S-phase population was observed in wild-type and MUS81^S87A^ cells. However, a higher percentage of BrdU-labelled G1 cells (G1*), a sign of a faster transit through the cell cycle, was observed in MUS81^S87D^ cells after 5 h of chase as compared with the wild-type ones (Fig. 6 B and C). As expected, in wild-type cells, inhibition of WEE1 (WEE1i) resulted in a faster progression from S-phase to mitosis, resulting in a strong increase of BrdU-positive cells in the subsequent G1-phase. In contrast, and interestingly, expression of the unphosphorylable S87A-MUS81 protein completely reverted the effect of WEE1i on cell cycle. Surprisingly, a reduction of the number of BrdU-positive G1 cells after WEE1 inhibition was also observed in cells expressing the phosphomimetic S87D-MUS81 mutant. Consistent results were also obtained by analysing progression of a BrdU-pulse labelled population to mitosis by anti-pS10H3/BrdU double immunofluorescence (Fig. 6 D). Indeed, treatment of wild-type cells with WEE1i greatly increased the number of BrdU-positive cells accumulated in mitosis by Noc, while expression of the S87A-MUS81 mutant reverted the phenotype. Of note, inhibition of WEE1 failed to increase the fraction of BrdU-positive mitosis detected in MUS81^S87D^ cells. Inhibition of CDK1 can prevent unscheduled activation of MUS81 by WEE1i (18). As shown in Figure S9 A, inhibition of CDK1 reverted the progression of BrdU-labelled S-phase cells to G2/M and the subsequent G1 that is stimulated by WEE1 inhibition, irrespective of the presence of a mutant MUS81 protein. Suppression of the WEE1i-induced premature mitotic entry is also obtained with EME2 or SLX4 depletion (18). In wild-type cells, depletion of EME2 reduced the unscheduled progression from S to G2/M and G1, which is stimulated by WEE1 inhibition, but only partially (Figure S9 B and C). Of note, depletion of EME2 reduced the limited progression from S to G2/M and G1 observed in the S87A-MUS81 mutant, while was largely ineffective in modulating the phenotype of the S87D-MUS81 mutant, either in the absence or in the presence of WEE1i. A consistent effect was observed when we analysed the S-M progression by anti-pS10H3/BrdU double immunofluorescence (Figure S9 D). Depletion of EME2 reduced the premature S-M transit induced by WEE1i in wild-type cells, while it minimally affects the MUS81^S87D^ phenotype. As expected, depletion of SLX4 completely reverts premature S-M transit independently on the deregulated S87 phosphorylation of MUS81 (Figure S9 D).

**Figure 6.**
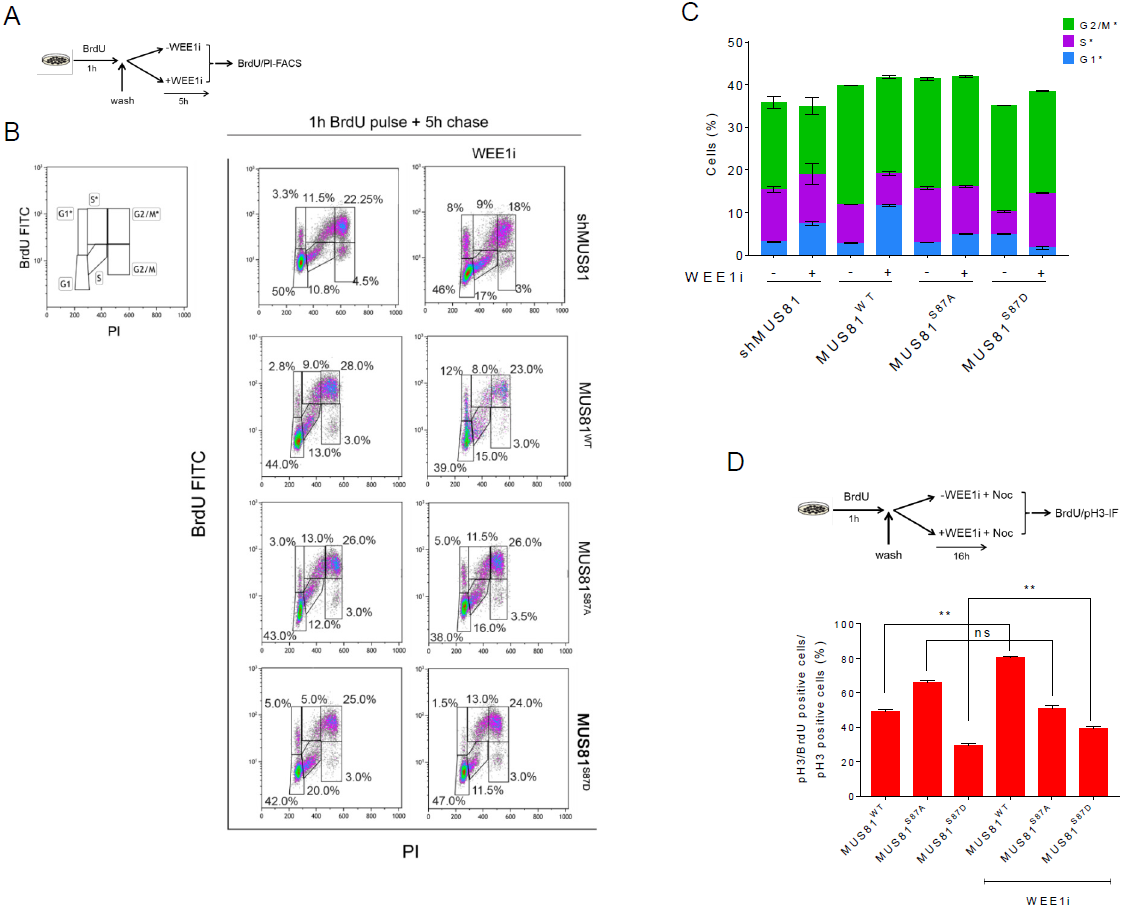
Premature mitotic entry mediated by unscheduled MUS81 function in S-phase depends on phosphorylation at S87. (A) Experimental workflow. (B) S-phase cells were pulse-labelled with BrdU and released in free medium for 5 h, in the presence or not of the WEE1 inhibitor MK-1775 (WEE1i). Progression of S-phase cells through the cell cycle was analysed by bivariate flow cytometry. The scheme indicates how each population was assigned. The star (*) denotes S-labelled, BrdU-positive, populations. Density plots depict mean BrdU intensities versus total nuclear DNA intensities (PI). The percentage of cells found in each phase of cell cycle is indicated. (C) The graph shows the percentage of cells in G2/M*, S* and G1* phase treated or not with WEE1i inhibitor MK-1775, relative to scatterplots in (B). (D) Immunofluorescence analysis of S-M progression. S-phase cells were labelled with a 1h BrdU pulse, and released in nocodazole to accumulate mitosis. Cells accumulated in mitosis for 16h, in presence or not of WEE1i, and immunostained using anti-BrdU and pS10H3 antibodies. Data are presented as mean ± standard error (SE) from three independent experiments. ** = p < 0.1; ns = not significant, ANOVA test.

These results indicate that phosphorylation of MUS81 by CK2 is absolutely required for the pathological S/M transit associated to WEE1 inhibition and deregulated phosphorylation of SLX4. Moreover, they suggest that the presence of a constitutively-active MUS81 complex and WEE1 inhibition does not synergize, but rather result in an apparent slow-down of the cell cycle.

### Regulated phosphorylation of MUS81 at Serine 87 is essential to prevent accumulation of genome instability

Downregulation of MUS81 induces mitotic defects and accumulation of bulky anaphase bridges as a consequence of poor resolution of replication/recombination intermediates prior to mitosis (19, 38). Our data suggest that S87 phosphorylation of MUS81 may affect function of the complex in mitosis. Hence, we analysed the presence of bulky anaphase bridges in cells expressing the wild-type MUS81 or the two S87 phosphomutants, exposed or not to a low-dose Aph (Fig. 7 A). Under unperturbed cell growth, the number of anaphase cells with bulky chromatin bridges was found elevated, albeit modestly, in both MUS81-depleted cells and in cells expressing the unphosphorylable S87A-MUS81 protein. In contrast, very few anaphases with bulky chromatin bridges were found in cells expressing the phosphomimic mutant of MUS81. In Aph-treated cells, the percentage of anaphases with bulky chromatin bridges was similar among cell lines, except un those expressing the phosphomimic S87D-MUS81 mutant, which showed less anaphase bridges. These data support the functional role of S87 phosphorylation of MUS81 in mitosis. Nevertheless, deregulation of S87 phosphorylation causes DNA damage in S-phase being involved in premature formation of the MUS81/EME1/SLX4 complex. Hence, we investigated whether expression of the S87 unphosphorylable or phosphomimic MUS81 mutant might undermine genome integrity. To this end, we analysed the number and type of chromosomal damage in metaphase spreads from cells treated or not with a low-dose Aph, which induces a mild replication stress that the MUS81complex contributes to fix. As shown in Fig. 7 B, mild replication stress increased the frequency of chromosome breakage in wild-type cells. As expected, the number of chromosome breaks detected on mild replication stress was slightly reduced upon MUS81 downregulation while, unexpectedly, it was only slightly enhanced by expression of the S87 unphosphorylable MUS81 mutant. In contrast, the frequency of chromosome breakage was significantly higher in cells expressing the S87D-MUS81 mutant already under unperturbed replication and increased further upon treatment with Aph. Interestingly, MUS81^S87D^ cells also showed complex chromosome aberrations, such as chromatid exchanges and pulverized metaphases, which were otherwise absent in wild-type or MUS81^S87A^ cells (Fig. 7 C). The unscheduled targeting of replication forks by the MUS81 endonuclease triggered by chemical inhibition of WEE1 is sufficient to induce chromosome pulverization (18). As we show that S87 phosphorylation predominates on WEE1 inhibition (Fig. 6), we evaluated if chromosome pulverization observed in cells treated with the WEE1i might be modulated by the presence of the two S87 phosphomutants of MUS81. As reported in Fig. 8 D, inhibition of WEE1 resulted in the appearance of pulverized metaphases in wild-type cells, and this phenotype greatly increased in response to Aph. As expected, the percentage of pulverisation was significantly reduced in MUS81-depleted cells respect to the wild-type, especially after Aph treatment (Fig. 8 D). Interestingly, WEE1i-dependent chromosome pulverisation was suppressed also by expression of the S87A-MUS81 mutant (Fig. 7 D). Conversely, MUS81^S87D^ cells showed chromosome pulverization even in absence of WEE1i, and the phenotype was significantly enhanced in its presence, suggesting that pharmacological override of cell cycle control and unscheduled targeting of replication forks by the MUS81/EME1 complex act synergistically. Interestingly, while depletion of EME2 partially rescued the pulverization phenotype associated to inhibition of WEE1 in wild-type cells it failed to modulate the pulverisation detected in MUS81^S87D^ cells (Figure S10 A and B), confirming that expression of the phosphomimetic MUS81 mutant preferentially engages the MUS81/EME1 complex.

**Figure 7.**
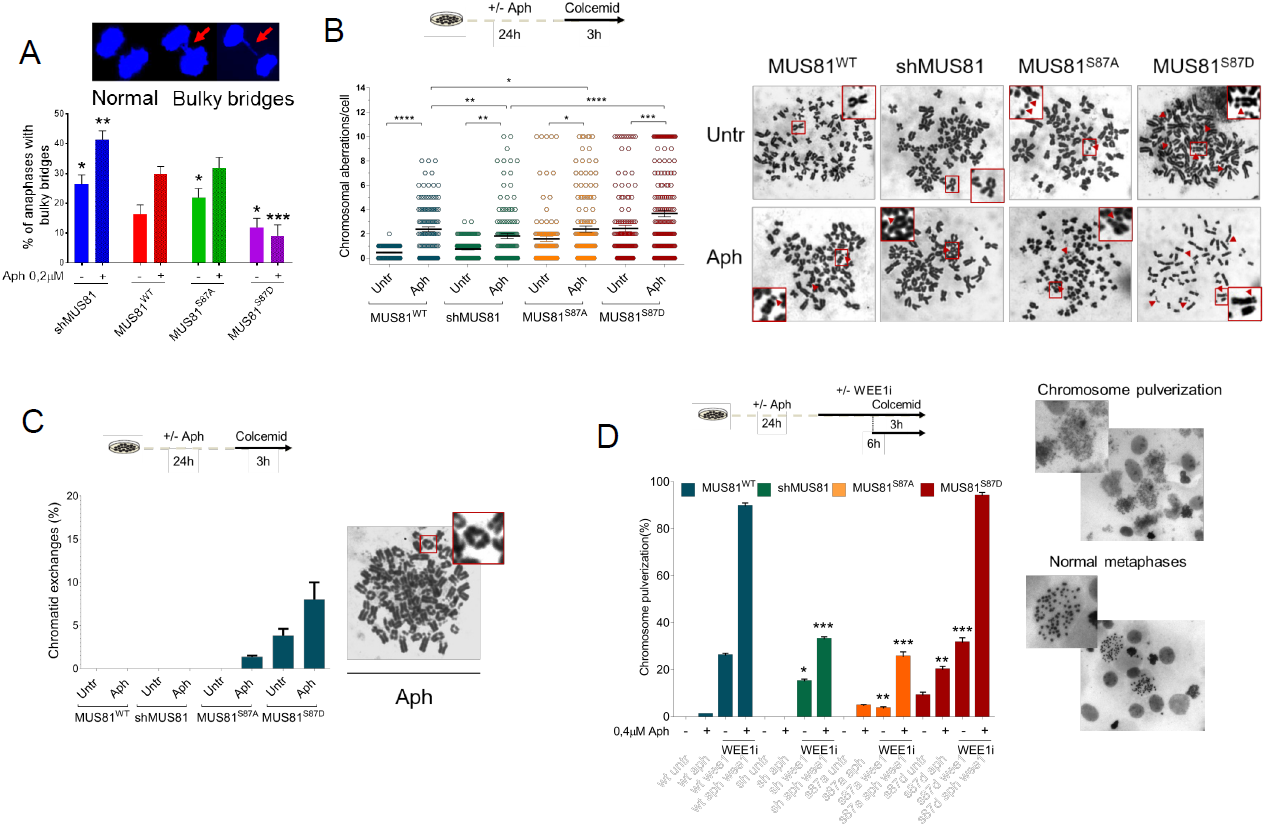
Regulated phosphorylation of MUS81 at S87 is essential to prevent accumulation of genome instability. (A) Analysis of the bulky anaphases bridges in MUS81 phosphomutants treated with a low-dose Aphidicolin (Aph). The graph shows the fractions of mitotic cells with bulky anaphase bridges. Error bars indicate SE; n = 3 (>50 mitotic cells were analyzed in each population). Data are presented as mean ± standard error (SE) from three independent experiments. * = p < 0.5; ** = p< 0.1; *** p<0.01, ANOVA test. Significance is reported compared to the wild-type. Representative images of single anaphases from the phosphomimetic MUS81 mutant are shown. Arrows indicate bridges. (B) Experimental scheme for evaluation of chromosomal aberrations is shown. Dot plot shows the number of chromosome aberrations per cell. Data are presented as means of three independent experiments. Horizontal black lines represent the mean ± SE. (ns, not significant; **, p < 0.01; ***, p < 0.001; ****, p < 0.0001 two-tailed Student’s t test). Representative Giemsa-stained metaphases are given. Arrows in red indicate chromosomal aberrations. (C) Analysis of the frequency of metaphase spreads with chromatid exchanges in cells treated and processed as in (B). Bar graph shows the percentage of chromatid exchanges per metaphase cell. Data are presented as means of three independent experiments. Horizontal black lines represent the mean ± SE. A representative Giemsa-stained metaphase with chromatid exchanges is given. (D) Experimental scheme for evaluation of chromosomal aberrations is shown. Bar graph shows the frequency of pulverized metaphases per metaphase cell. Data are presented as means of three independent experiments. Error bars representing standard errors are not shown or clarity but are < 15 of the mean (*, p<0.5 **, p < 0.01; ***, p < 0.001; two-tailed Student’s t test). Representative Giemsa-stained metaphases are given for both normal and pulverized phenotype. Significance is reported compared to the wild-type.

**Figure 8.**
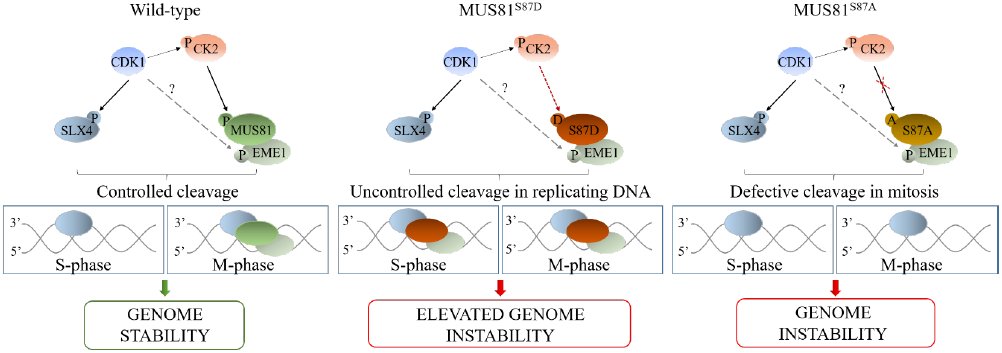
Summarizing model to illustrate the contribution of CK2-dependent phosphorylation to the MUS81/EME1/SLX4 regulation. See discussion for details.

Therefore, we conclude that phosphorylation of MUS81 S87 by CK2 is required to support function of the MUS81 complex during resolution of intermediates accumulating under mild replication stress, but also that loss of regulated phosphorylation strongly undermines genome integrity.

## DISCUSSION

The structure-specific endonuclease MUS81/EME1 plays important roles in the resolution of recombination intermediates, however, its function needs to be carefully regulated to avoid an unscheduled targeting intermediates during DNA replication, which may result in genome instability. Regulation of the MUS81 complex has been mostly investigated in yeast, and several publications demonstrated that cell cycle-dependent phosphorylation of the Mms4^EME1^ subunit by Cdc28^CDK1^ or Cdc5^PLK1^ ensures activation of the MUS81 complex in G2/M (16, 39, 40). In yeast, the MUS81 complex can be also regulated by checkpoint kinases in response to DNA damage (16, 17). In human cells, a cell cycle-dependent regulation of the MUS81 complex has been indirectly inferred from association with PLK1, and the presence of phosphorylated isoforms of EME1 in mitosis (41, 42). The functional role of such events and the identity of the targeted residues are almost unknown, and only recently a mechanistic link between CDK1-dependent phosphorylation and activation of the human SLX4-MUS81-EME2 complex has been revealed (18). However, in most cases, the phenotype associated with loss of MUS81 complex regulation has been deducted from inhibition of cell cycle-related kinases, such as CDK1 or PLK1, so that the observed effect may derive also from perturbation of cell cycle progression *per se* or from altered function of other targets.

Here, we find that the biological function of the MUS81-EME1 complex in human cells is positively regulated by CK2, which phosphorylates Serine 87 (S87) of the MUS81 subunit in mitosis. Interestingly, phosphorylation at S87 increases from late G2 to prophase and disappears as soon as cells proceed to metaphase. Activation of the MUS81-EME1 complex during prophase or pro-metaphase has been hypothesised from interaction with its key partners using synchronised cells (18). Our data on S87 phosphorylation confirm that the human MUS81-EME1 complex becomes active in late G2 and early mitosis and suggest that pS87MUS81 is excluded from chromatin in metaphase. Our findings also show that MUS81 S87 phosphorylation is very low or undetectable in S-phase. The MUS81 complex(es) needs to be switched off during normal replication (4), and downregulation of S87 phosphorylation is consistent with the need to avoid adventitious targeting of replication intermediates. Although MUS81 can form distinct heterodimeric complexes with EME1 or EME2 (20), our data show that phosphorylation of S87 is mainly detected in the MUS81-EME1 complex. This is consistent with the MUS81-EME1 dimer being active in mitosis and suggests that different phosphorylation events may be involved in the regulation of specific MUS81 complexes. From this point of view, phosphorylation of the invariant subunit of the MUS81 heterodimers could be advantageous to direct association of MUS81 with EME1 or EME2, and to modulate endonucleolytic cleavage *in vivo*. From this point of view, although the amount of MUS81 in chromatin is unchanged by the S87 phosphorylation status, the different level of chromatin-associated EME1 or EME2 in the S87 phosphorylation mutants of MUS81 may reflect the ability to form different complexes.

Experiments using inhibitors of cell cycle kinases provided clues about consequences of unscheduled activation of the MUS81 complex on viability and chromosome stability in human cells (18, 19, 32). Our findings clearly demonstrate that expression of a phosphomimic S87D-MUS81 mutant is detrimental to cell proliferation because of the generation of DSBs in S-phase cells. Interestingly, such unscheduled formation of DSBs in S-phase is largely prevented by ectopic expression of the bacterial RuvA protein. RuvA is a Holliday junction-binding protein, and its ectopic expression in human cells counteracts generation of DSBs by another structure-specific endonuclease, GEN1 (37). Hence, from a mechanistic point of view, a mutation mimicking constitutive phosphorylation of MUS81 at S87 is sufficient to unleash targeting of HJ-like intermediates arising during normal replication and this is linked to generation of DNA damage. Much more interestingly, under unperturbed cell growth, expression of the S87D-MUS81 mutant results in a chromosome fragility phenotype, which is more marked than that observed in cells expressing the unphosphorylable S87A-MUS81 form. In addition, cells expressing the phosphomimic S87D-MUS81 mutant not only have elevated number of chromosome breaks and gaps, but also have a striking accumulation of radial chromosomes. Since the presence of radial chromosomes is associated with repair of DSBs at collapsed forks by NHEJ (43), unscheduled formation of DSBs by MUS81 complex in S-phase may engage end-joining repair in addition to homologous recombination, which may promote gross chromosomal rearrangements.

In unperturbed cells, loss of the MUS81 complex function results in mitotic defects including accumulation of bulky anaphase bridges (19, 38). We show that cells expressing the S87A-MUS81 mutant recapitulate the high number of bulky anaphase bridges observed in cells depleted of MUS81, although they show few chromosome breaks in metaphase. In contrast, expression of the phosphomimic S87D-MUS81 results in few bulky anaphase bridges, but in a striking chromosome fragility. Hence, formation of DSBs by an unscheduled function of the MUS81 complex during a normal S-phase is expected to be much more detrimental to genome stability of that associated with activation of the MUS81 complex at collapsed replication forks, while loss of resolution activity in early mitosis is better sustained, probably because of backup activities (7, 9, 11).

Prolonged or pathological replication fork arrest has been associated to induction of MUS81-dependent DSBs (7, 8, 11, 44). Our data show that MUS81 S87 phosphorylation is low in cells treated with HU for 24 h, a condition known to promote MUS81 function at collapsed replication forks (8), indicating distinct regulatory mechanisms of the human MUS81 complex during mild or persisting replication stress. However, we also provide evidence that CK2 inhibition reverts formation of MUS81-dependent DSBs after HU treatment. Hence, CK2-dependent phosphorylation on other residues of MUS81 or EME1/2 may be involved in regulating the MUS81 complex under different conditions, as reported in yeast (16, 17). Alternatively, CK2 may target other proteins required for MUS81 complex function under conditions of persisting replication stress. Interestingly, CK2 phosphorylates, among others, the RAD51 recombinase (25). Of note, RAD51 or RAD52 are involved in the formation of the MUS81 complex substrates at demised replication forks (7, 44). Clarifying this interesting point clearly goes beyond the scope of this work and deserves future investigations.

In contrast to persisting or pathological replication stress, a mild condition of replication perturbation that does not arrest cells in S-phase can stimulate S87 phosphorylation of MUS81. Such mild condition of replication stress has been reported to trigger the function of the MUS81-EME1 complex to resolve intermediates at under-replicated regions, such as common fragile sites (12, 13). Our data on the cell cycle-specificity of MUS81 S87 phosphorylation suggest that this function of the MUS81 complex occurs post-replication or in mitosis when common fragile sites loci might conclude their replication (29). Strikingly, expression of the unphosphorylable MUS81 S87A mutant enhances chromosomal damage in cells treated with low-dose Aph only slightly, while the presence of a phosphomimic MUS81 mutant dramatically increases the amount of chromosome breakage and radial chromosomes after a mild replication stress.

Hence, as in untreated cells, the phenotype deriving from unscheduled targeting of perturbed replication forks by the MUS81 complex predominates over that resulting from loss of function in G2/M.

Our experiments indicate that phosphorylation of MUS81 S87 is important to establish a correct interaction between MUS81 and SLX4. SLX4 is a versatile scaffold involved in recruitment of multiple endonucleases (42, 45). In particular, activation of MUS81/EME1 in mitotic cells requires association with SLX4 through a region localized in the first 90 aminoacids of MUS81 (18, 33, 41). Interestingly, deletion of the SLX4-interacting region of MUS81 broadens substrate specificity, allowing the complex to target much more easily also replication intermediates at perturbed forks (46). Serine 87 is within the region interacting with SLX4 and its phosphorylation makes MUS81 more prone to associate with SLX4, providing a mechanistic explanation to the unscheduled targeting by the MUS81 S87D protein of HJ-like and possibly other branched intermediates during replication.

It has been recently reported that inhibition of WEE1 licences association of MUS81/EME2 with SLX4 during S-phase resulting in a wide chromosome instability in the form of pulverized metaphases (18). Interestingly, the dramatic effect of WEE1 inhibition on premature cell cycle progression and chromosome pulverization is completely prevented by the unphosphorylable MUS81 S87A mutant, reinforcing the strong functional value of S87 phosphorylation. In our experimental conditions, however, depletion of EME2 substantially reduces but not suppresses WEE1i-associaated phenotypes. In particular, expression of the S87D-MUS81 mutant induces a phenotype that is only minimally affected by EME2 depletion. As S87 phosphorylation occurs mainly in the MUS81/EME1 complex, it is conceivable that expression of the phosphomimic protein favours engagement of the MUS81/EME1 complex while, in wild-type cells, both complexes might contribute to the WEE1i-dependent phenotypes. Alternatively, the relative amount of each MUS81 complex may be cell-specific and affect also the genetic dependency of the WEE1i-dependent effects. Moreover, as WEE1-mediated promotion of MUS81/SLX4 interaction involves CDK1-dependent phosphorylation of SLX4 (18), our data indicate that both the partners must be modified to promote a productive interaction. CK2 is a crucial kinase in mitosis, and is positively regulated by CDK1 (22). From this point of view, human cells might have evolved a redundant regulatory mechanism to restrain MUS81 complex activity in late G2 and M phase. Indeed, elevated CDK1 level may directly contribute to activate the MUS81 complex phosphorylating EME1(2), while, indirectly, may enhance activity of CK2 that, in turn, targets MUS81 licensing interaction with an already phosphorylated SLX4 (Fig. 8). This elaborate regulatory mechanism will allow cleavage of branched DNA intermediates during a narrow window at the beginning of mitosis, contributing to limit chromosome instability.

Our results may also have strong implications for the onset of genome instability in cancer. Indeed, CK2 has been found overexpressed in many human tumours (47). Hence, it is tempting to speculate that in CK2-overexpressing cancer cells also the biological function of MUS81/EME1 may be elevated and contribute strongly to genome instability and, possibly, aggressiveness. Further studies will be needed to evaluate the status of MUS81 S87 in human tumours together with level and type of genome instability, in order to see if there is any correlation with the expression of CK2.

Altogether, our study provides the first mechanistic insight into the regulation of the human MUS81 complex, with functional implications on the consequences of deregulated processing of replication intermediates during unperturbed or minimally-perturbed DNA replication.

## ACKNOWLEDGMENTS

We are grateful to Dr. Agata Smogorzewska (Rockefeller University, New York, USA) for the FA-P cells. The pEF1a-IRES-Neo was a gift from Thomas Zwaka (Addgene plasmid #28019). We thank Drs. Achille Pellicioli and Federica Marini for helpful discussion. This work was supported by investigator grants from Associazione Italiana per la Ricerca sul Cancro (AIRC) to P.P. (IG#13398 and IG#17383) and to A.F. (IG#15410), and by Nando-Peretti Foundation to P.P. (grant n. 2012-113).

## AUTHORS CONTRIBUTION

A.P. performed protein interaction experiments, characterization of S87 phosphorylation and contributed to phenotypic experiments. G.M.P. characterized the MUS81 mutants, performed DNA damage and flow cytometry experiments, and studied on mitotic phenotypes. I.M. performed in vitro kinase assays and generated MUS81 mutants. M.S. contributed to design and analyse flow cytometry experiments. A.M. performed analysis of endonuclease assays. F.M. contributed to design and supervised endonuclease assays. V.M. performed analysis of chromosomal damage. S.R. performed MS/MS. E.M. performed chromatin fractionation experiments. L.Z. supervised MS/MS experiments. P.P and A.F. designed experiments and analysed data. All authors contributed to design experiments. P.P. and A.F. wrote the paper. All authors contributed to revise the paper.

## REFERENCES

1. Sarbajna, S. and West, S.C. (2014) Holliday junction processing enzymes as guardians of genome stability. Trends Biochem. Sci., 39, 409–419.

2. Mankouri, H.W., Huttner, D. and Hickson, I.D. (2013) How unfinished business from S-phase affects mitosis and beyond. EMBO J., 32, 2661–71.

3. Rass, U. (2013) Resolving branched DNA intermediates with structure-specific nucleases during replication in eukaryotes. Chromosoma, 122, 499–515.

4. Matos, J. and West, S.C. (2014) Holliday junction resolution: Regulation in space and time. DNA Repair (Amst)., 19, 176–181.

5. Ashton, T.M., Mankouri, H.W., Heidenblut, A., McHugh, P.J. and Hickson, I.D. (2011) Pathways for Holliday junction processing during homologous recombination in Saccharomyces cerevisiae. Mol. Cell. Biol., 31, 1921–1933.

6. Osman, F. and Whitby, M.C. (2007) Exploring the roles of Mus81-Eme1/Mms4 at perturbed replication forks. DNA Repair (Amst)., 6, 1004–1017.

7. Murfuni, I., Basile, G., Subramanyam, S., Malacaria, E., Bignami, M., Spies, M., Franchitto, A. and Pichierri, P. (2013) Survival of the Replication Checkpoint Deficient Cells Requires MUS81- RAD52 Function. PLoS Genet., 9, e1003910.

8. Hanada, K., Budzowska, M., Davies, S.L., van Drunen, E., Onizawa, H., Beverloo, H.B., Maas, A., Essers, J., Hickson, I.D. and Kanaar, R. (2007) The structure-specific endonuclease Mus81 contributes to replication restart by generating double-strand DNA breaks. Nat. Struct. Mol. Biol., 14, 1096–1104.

9. Franchitto, A., Pirzio, L.M., Prosperi, E., Sapora, O., Bignami, M. and Pichierri, P. (2008) Replication fork stalling in WRN-deficient cells is overcome by prompt activation of a MUS81-dependent pathway. J. Cell Biol., 183, 241–252.

10. Froget, B., Blaisonneau, J., Lambert, S. and Baldacci, G. (2008) Cleavage of stalled forks by fission yeast Mus81/Eme1 in absence of DNA replication checkpoint. Mol. Biol. Cell, 19, 445–56.

11. Murfuni, I., Nicolai, S., Baldari, S., Crescenzi, M., Bignami, M., Franchitto, a and Pichierri, P. (2012) The WRN and MUS81 proteins limit cell death and genome instability following oncogene activation. Oncogene, 32, 610–20.

12. Naim, V., Wilhelm, T., Debatisse, M. and Rosselli, F. (2013) ERCC1 and MUS81-EME1 promote sister chromatid separation by processing late replication intermediates at common fragile sites during mitosis. Nat. Cell Biol., 15, 1008–15.

13. Ying, S., Minocherhomji, S., Chan, K.L., Palmai-Pallag, T., Chu, W.K., Wass, T., Mankouri, H.W., Liu, Y. and Hickson, I.D. (2013) MUS81 promotes common fragile site expression. Nat. Cell Biol., 15, 1001–7.

14. Wild, P. and Matos, J. (2016) Cell cycle control of DNA joint molecule resolution. Curr. Opin. Cell Biol., 40, 74–80.

15. Princz, L.N., Wild, P., Bittmann, J., Aguado, F.J., Blanco, M.G., Matos, J. and Pfander, B. (2017) Dbf4-dependent kinase and the Rtt107 scaffold promote Mus81-Mms4 resolvase activation during mitosis. EMBO J., 10.15252/embj.201694831.

16. Dehé, P.-M., Coulon, S., Scaglione, S., Shanahan, P., Takedachi, A., Wohlschlegel, J. a, Yates, J.R., Llorente, B., Russell, P. and Gaillard, P.-H.L. (2013) Regulation of Mus81-Eme1 Holliday junction resolvase in response to DNA damage. Nat. Struct. Mol. Biol., 20, 598–603.

17. Kai, M., Boddy, M.N., Russell, P. and Wang, T.S.F. (2005) Replication checkpoint kinase Cds1 regulates Mus81 to preserve genome integrity during replication stress. Genes Dev., 19, 919–932.

18. Duda, H., Arter, M., Gloggnitzer, J., Teloni, F., Wild, P., Blanco, M.G., Altmeyer, M. and Matos, J. (2016) A Mechanism for Controlled Breakage of Under-replicated Chromosomes during Mitosis. Dev. Cell, 39, 740–755.

19. Matos, J., Blanco, M.G. and West, S.C. (2013) Cell-cycle kinases coordinate the resolution of recombination intermediates with chromosome segregation. Cell Rep., 4, 76–86.

20. Pepe, A. and West, S.C. (2014) MUS81-EME2 promotes replication fork restart. Cell Rep., 7, 1048–1055.

21. Fekairi, S., Scaglione, S., Chahwan, C., Taylor, E.R., Tissier, A., Coulon, S., Dong, M.Q., Ruse, C., Yates, J.R., Russell, P., et al. (2009) Human SLX4 Is a Holliday Junction Resolvase Subunit that Binds Multiple DNA Repair/Recombination Endonucleases. Cell, 138, 78–89.

22. Meggio, F. and Pinna, L. a (2003) One-thousand-and-one substrates of protein kinase CK2? FASEB J., 17, 349–368.

23. Franchin, C., Borgo, C., Zaramella, S., Cesaro, L., Arrigoni, G., Salvi, M. and Pinna, L.A. (2017) Exploring the CK2 Paradox: Restless, Dangerous, Dispensable. Pharmaceuticals (Basel)., 10, 11.

24. Spycher, C., Miller, E.S., Townsend, K., Pavic, L., Morrice, N. a., Janscak, P., Stewart, G.S. and Stucki, M. (2008) Constitutive phosphorylation of MDC1 physically links the MRE11-RAD50- NBS1 complex to damaged chromatin. J. Cell Biol., 181, 227–40.

25. Yata, K., Lloyd, J., Maslen, S., Bleuyard, J.-Y., Skehel, M., Smerdon, S.J. and Esashi, F. (2012) Plk1 and CK2 Act in Concert to Regulate Rad51 during DNA Double Strand Break Repair. Mol. Cell, 45, 371–383.

26. Kim, S.T. (2005) Protein kinase CK2 interacts with Chk2 and phosphorylates Mre11 on serine 649. Biochem. Biophys. Res. Commun., 331, 247–252.

27. Chapman, J.R. and Jackson, S.P. (2008) Phospho-dependent interactions between NBS1 and MDC1 mediate chromatin retention of the MRN complex at sites of DNA damage. EMBO Rep., 9, 795–801.

28. Ammazzalorso, F., Pirzio, L.M., Bignami, M., Franchitto, A. and Pichierri, P. (2010) ATR and ATM differently regulate WRN to prevent DSBs at stalled replication forks and promote replication fork recovery. EMBO J., 29, 3156–3169.

29. Minocherhomji, S., Ying, S., Bjerregaard, V.A., Bursomanno, S., Aleliunaite, A., Wu, W., Mankouri, H.W., Shen, H., Liu, Y. and Hickson, I.D. (2015) Replication stress activates DNA repair synthesis in mitosis. Nature, 528, 286–290.

30. Pichierri, P., Franchitto, a, Mosesso, P., Proietti de Santis, L., Balajee, a.. and Palitti, F. (2000) Werner’s syndrome lymphoblastoid cells are hypersensitive to topoisomerase II inhibitors in the G2 phase of the cell cycle. Mutat. Res. Repair, 459, 123–133.

31. Dehé, P.-M. and Gaillard, P.-H.L. (2017) Control of structure-specific endonucleases to maintain genome stability. Nat. Rev. Mol. Cell Biol., 18, 315–330.

32. Forment, J. V., Blasius, M., Guerini, I. and Jackson, S.P. (2011) Structure-specific DNA endonuclease mus81/eme1 generates DNA damage caused by chk1 inactivation. PLoS One, 6, e23517.

33. Nair, N., Castor, D., Macartney, T. and Rouse, J. (2014) Identification and characterization of MUS81 point mutations that abolish interaction with the SLX4 scaffold protein. DNA Repair (Amst)., 24, 131–137.

34. Pfander, B. and Matos, J. (2017) Control of Mus81 nuclease during the cell cycle. FEBS Lett., 591, 2048–2056.

35. Fugger, K., Chu, W.K., Haahr, P., Kousholt, A.N., Beck, H., Payne, M.J., Hanada, K., Hickson, I.D. and Sørensen, C.S. (2013) FBH1 co-operates with MUS81 in inducing DNA double-strand breaks and cell death following replication stress. Nat. Commun., 4, 1423.

36. Ciccia, A., Ling, C., Coulthard, R., Yan, Z., Xue, Y., Meetei, A.R., Laghmani, E.H., Joenje, H., McDonald, N., de Winter, J.P., et al. (2007) Identification of FAAP24, a Fanconi Anemia Core Complex Protein that Interacts with FANCM. Mol. Cell, 25, 331–343.

37. Malacaria, E., Franchitto, A. and Pichierri, P. (2017) SLX4 Prevents GEN1-Dependent DSBs During DNA Replication Arrest Under Pathological Conditions in Human Cells. Sci. Rep., 7, 44464.

38. Garner, E., Kim, Y., Lach, F.P., Kottemann, M.C. and Smogorzewska, A. (2013) Human GEN1 and the SLX4-Associated Nucleases MUS81 and SLX1 Are Essential for the Resolution of Replication-Induced Holliday Junctions. Cell Rep., 5, 207–215.

39. Matos, J., Blanco, M.G., Maslen, S., Skehel, J.M. and West, S.C. (2011) Regulatory control of the resolution of DNA recombination intermediates during meiosis and mitosis. Cell, 147, 158–172.

40. Gallo-Fernández, M., Saugar, I., Ortiz-Bazán, M.Á., Vázquez, M.V. and Tercero, J.A. (2012) Cell cycle-dependent regulation of the nuclease activity of Mus81-Eme1/Mms4. Nucleic Acids Res., 40, 8325–8335.

41. Wyatt, H.D.M., Sarbajna, S., Matos, J. and West, S.C. (2013) Coordinated actions of SLX1-SLX4 and MUS81-EME1 for holliday junction resolution in human cells. Mol. Cell, 52, 234–247.

42. Svendsen, J.M., Smogorzewska, A., Sowa, M.E., O’Connell, B.C., Gygi, S.P., Elledge, S.J. and Harper, J.W. (2009) Mammalian BTBD12/SLX4 Assembles A Holliday Junction Resolvase and Is Required for DNA Repair. Cell, 138, 63–77.

43. Kasparek, T.R. and Humphrey, T.C. (2011) DNA double-strand break repair pathways, chromosomal rearrangements and cancer. Semin. Cell Dev. Biol., 22, 886–97.

44. Hengel, S.R., Malacaria, E., Folly da Silva Constantino, L., Bain, F.E., Diaz, A., Koch, B.G., Yu, L., Wu, M., Pichierri, P., Spies, M.A., et al. (2016) Small-molecule inhibitors identify the RAD52- ssDNA interaction as critical for recovery from replication stress and for survival of BRCA2 deficient cells. Elife, 5.

45. Muñoz, I.M., Hain, K., Déclais, A.-C., Gardiner, M., Toh, G.W., Sanchez-Pulido, L., Heuckmann, J.M., Toth, R., Macartney, T., Eppink, B., et al. (2009) Coordination of structure-specific nucleases by human SLX4/BTBD12 is required for DNA repair. Mol. Cell, 35, 116–27.

46. Wyatt, H.D.M., Laister, R.C., Martin, S.R., Arrowsmith, C.H. and West, S.C. (2017) The SMX DNA Repair Tri-nuclease. Mol. Cell, 65, 848–860.e11.

47. Ruzzene, M. and Pinna, L.A. (2010) Addiction to protein kinase CK2: a common denominator of diverse cancer cells? Biochim. Biophys. Acta, 1804, 499–504.

